# Mutation rate variability in viral populations: implications for lethal mutagenesis

**DOI:** 10.1101/2025.05.16.654520

**Authors:** Sarah Arcos, Adam S. Lauring

**Author notes:** Sarah Arcos **Email:**. **Author Contributions:** S.A. designed research, performed research, analyzed the data, and wrote the paper. A.S.L. designed research and edited the paper. **Competing Interest Statement:** No competing interests.

## Abstract

Lethal mutagenesis is a strategy to achieve viral extinction by drugging viral mutation rates beyond an extinction threshold. Accurate estimation of the extinction threshold is critical, as elevating viral mutation rates near, but not past this threshold increases the likelihood of mutations that could result in drug resistance, vaccine escape, or increased pathogenesis. Traditional models of lethal mutagenesis rely on the Poisson distribution, which assumes a uniform mutation rate across individuals. Yet, RNA viruses like influenza A virus (IAV) can have varied mutation rates due to mutations in the polymerase complex. This variability suggests that lethal mutagenesis models incorporating mutation rate diversity, such as ones using the gamma-Poisson distribution, may be more accurate for RNA viruses. Poisson models assume count data have equal mean and variance, while gamma-Poisson counts are overdispersed (variance greater than mean). Here we provide experimental data showing that IAV mutations are overdispersed, indicating that the gamma-Poisson distribution is more appropriate for modeling IAV mutations. Modeling of lethal mutagenesis using the gamma-Poisson distribution reveals that the degree of overdispersion is critical in determining survival or extinction. Increased overdispersion shifts the extinction threshold higher, indicating that Poisson-based models have underestimated the mutation rate required to achieve viral extinction and avoid viral escape or accelerated evolution. Furthermore, time to extinction in simulated populations is significantly longer with gamma-Poisson-based models than Poisson-based. This investigation of how mutation rate variability affects lethal mutagenesis will directly impact antiviral drug design and strategy, thus advancing efforts to combat virus outbreaks and future pandemics.

**Significance Statement:** RNA viruses mutate at high rates. Some antiviral drugs work by further increasing these rates to unsustainable levels, a process called lethal mutagenesis. In this work, we discover that mutation rates vary significantly throughout viral populations. This discovery changes our understanding of how mutations and mutation rates should be modeled. By considering variability in mutation rates, our new model reveals that lethal mutagenesis is less effective as an antiviral drug strategy than previously thought.

## Introduction

In 1943, Luria and Delbrück proved that mutations in bacteria are random through a series of groundbreaking experiments (1). In their work, they proposed that the distribution of mutations across individuals in a bacterial population would follow a Poisson distribution. Over the past 80 years, this assumption of Poisson-distributed mutations has been applied to every kingdom of life and used to estimate species mutation rates. For a Poisson-based model to be an accurate reflection of the biology mutations must be 1) random and independent and 2) occur according to a *fixed* rate. While generally valid in most systems, the first assumption may not be accurate in all biological contexts. For example, specific mutation types can be induced by antiviral proteins like APOBEC. However, the second assumption has not been seriously examined, despite evidence that it may be substantially inaccurate, particularly for RNA viruses or other organisms with high mutation rates.

Accurate modeling of viral mutation rates is critical for the successful use of mutagenic antiviral drugs. These drugs are thought to work by *lethal mutagenesis* – a process of viral extinction through drugging mutation rates to unsustainable levels (2–8). Current models of lethal mutagenesis assume that viral mutations follow a Poisson distribution, in accordance with Luria and Delbrück’s earlier work. As most mutations are deleterious, an increase in mutations leads to a decrease in the fitness of offspring. When the number of surviving offspring is less than the number of parents, a viral population will eventually go extinct. This theory of lethal mutagenesis is distinct from the concept of “error catastrophe”, which instead describes changes in the genotypic landscape of a population and can actually delay or prevent extinction (4). Lethal mutagenesis can be achieved by sufficiently drugging a viral population across an *extinction threshold*, which is determined in the simplest model by viral mutation rate and fecundity (i.e., the number of viable offspring per individual) (Figure 1, Equation 1) (4). However, if a viral population is treated with a mutagenic drug, but the extinction threshold is not reached, the population will produce increased mutations without going extinct. Thus, the consequences of mutagenic drug treatments that do not meet the extinction threshold are dire, including increased risk for antiviral drug resistance and antibody escape, expansion to new hosts or cell types, and increased virulence. In fact, recent work has shown that sub-lethal treatment of SARS-CoV-2 in human patients with the antiviral drug molnupiravir led to transmission of persistent mutagenic signatures (9).

**Figure 1.**
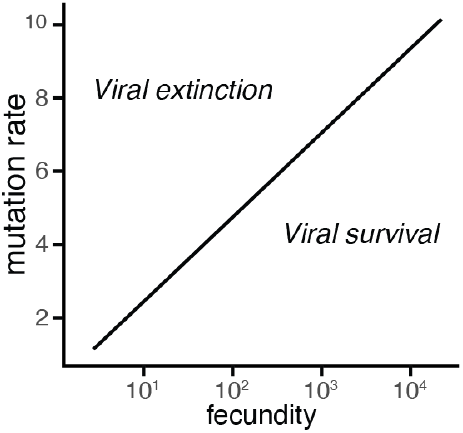
Diagram of the extinction threshold for RNA viruses, according to Equation 1, adapted from (4). Viruses with mutation rate and fecundity that fall below the threshold will survive, while viruses above the threshold will go extinct. Mutation rate is the mutations per replicated genome; fecundity is the number of offspring per individual.

In this work, we critically examine the assumption that mutations occur according to a *fixed* chance in Influenza A viruses (IAVs). We find that mutation rates within an IAV population are variable and that mutations are overdispersed on a genomic level. This variability means that viral mutation counts are more accurately modeled using the gamma-Poisson distribution, a statistical distribution that can account for *variable* rates. We then use the gamma-Poisson distribution to refine models of lethal mutagenesis, revealing that the effectiveness of mutagenic drug treatment is dependent upon the amount of mutation rate variation within viral populations. Lastly, we demonstrate that Poisson-based models of the lethal mutagenesis have systematically underestimated the extinction threshold, thus increasing the public health risks associated with current mutagenic drug treatment strategies.

## Results

### Evidence for mutation rate variability

Mutation rates among individuals within viral populations can vary due to environmental, stochastic, or genetic causes. There are many environmental factors that can affect mutation rates, including temperature, pH, and cell type or physiological state. However, for some RNA viruses, mutation rate appears to remain stable across physiological conditions and cell types (10, 11). Mutation rate may vary due to stochastic fluctuations in host cell composition, gene expression, or cell cycle state. Lastly, the high mutation rate of RNA viruses suggests that genetic changes substantially contribute to mutation rate variability. Based on our own empirical estimates of the IAV mutation rate, we estimate that there are on average 1-2 nonsynonymous mutations per polymerase complex per replicated genome (11). This suggests that RNA virus mutation rates are high enough to generate significant diversity in the biochemical structure and properties (including mutation rate) of an RNA virus polymerase within a single generation (Figure 2A). We can further assume the presence of genetic mutation rate variability within viral populations given that mutation rates themselves can evolve (5, 12–14); for evolution to occur, there must be a diversity of mutation rates upon which natural selection can act.

**Figure 2.**
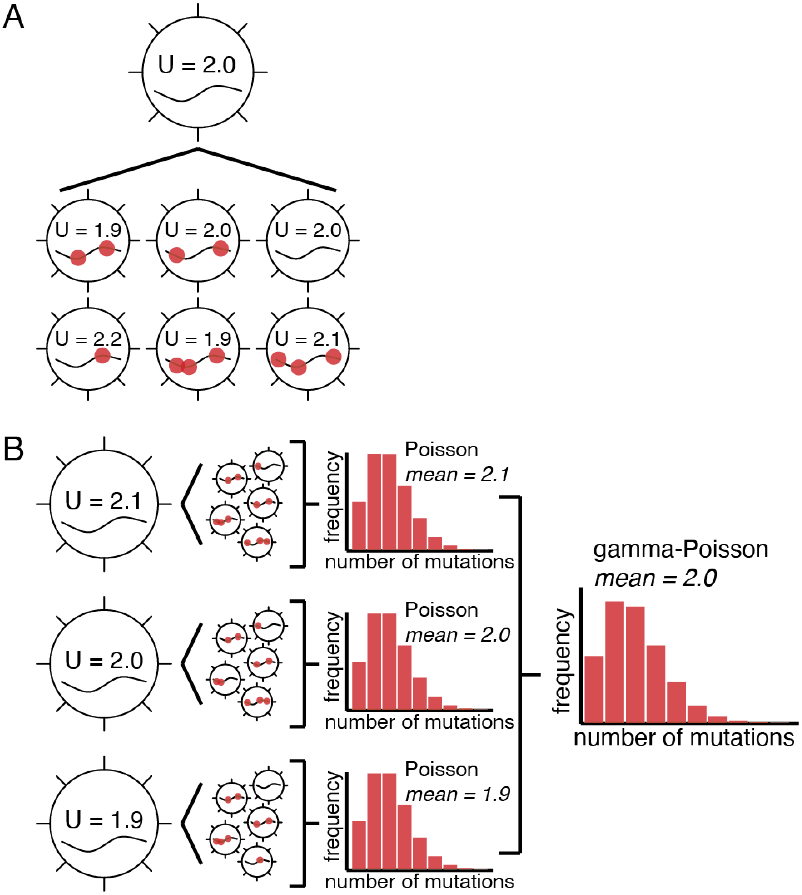
Viral populations contain diverse mutation rates. A) A single infectious unit can produce viral progeny with varying mutation rates due to mutations in the viral polymerase. Red dot indicates a mutation. Based on estimates of the viral mutation rate of IAV, 1-2 nonsynonymous mutations occur within the polymerase complex per replicated genome. B) A mixture of viral mutation rates within a population gives rise to mutations that follow a gamma-Poisson distribution (a mixture of Poisson distributions with gamma-distributed mutation rates). U, mutations per replicated genome.

If mutation rates indeed vary within viral populations, then the distribution of the number of mutations per genome per replication will not follow a Poisson distribution, as previously assumed (1), because it violates the fundamental Poisson assumption that the random events (mutations) occur according to a small, *fixed* chance. In the case of populations with high mutation rates, the chance of mutation is likely not *fixed*, but *variable* across the viral population. Therefore, a statistical distribution that models random events with rate variability should more accurately capture the true behavior of an RNA virus population. One such distribution is the gamma-Poisson distribution, which is a mixture of Poisson distributions with gamma-distributed rates (Figure 2B). This distribution is an appealing alternative because the gamma distribution is already used to model substitution rates in phylogenetic analyses, and mutation rates can reasonably be expected to be distributed similarly to substitution rates (15–17). Furthermore, the gamma-Poisson distribution was recently shown to be a better fit than the Poisson for mutation counts in a yeast model of cancer cells with mutation rate variability (18).

A key distinction between Poisson and gamma-Poisson distributions is that the mean and variance of a Poisson-distributed variable will be equal, while a gamma-Poisson-distributed variable can be *overdispersed* with a variance is greater than the mean. Therefore, the hypothesis that mutation rates are variable within viral populations can be tested empirically; if observed counts of mutations per genome per replication are overdispersed, then this indicates that the underlying population contains variable mutation rates (Figure 2B). While this variability could be due to a combination of environmental, stochastic, and genetic causes discussed above, we can focus on the genetic component by using a controlled cell culture environment. To test this hypothesis, we performed mutation accumulation experiments on Influenza A/Netherlands/499/2017(H3N2) over a single round of replication (Figure 3). We propagated the virus for a single round of replication (Figure 3A) and then performed endpoint dilution to isolate infectious viruses and their genomes. These infectious clones were sequenced to identify and count *fixed* (>95% frequency) mutations (see Methods). We observed that mutation counts among viral clones after a single round of IAV replication are indeed overdispersed, with an index of dispersion (ratio of variance to mean) of 1.16 (Figure 3B, left panel). This finding is consistent with previous work showing that individual vesicular stomatitis virus progeny isolated from single cells have variable spontaneous mutation rates (19).

**Figure 3.**
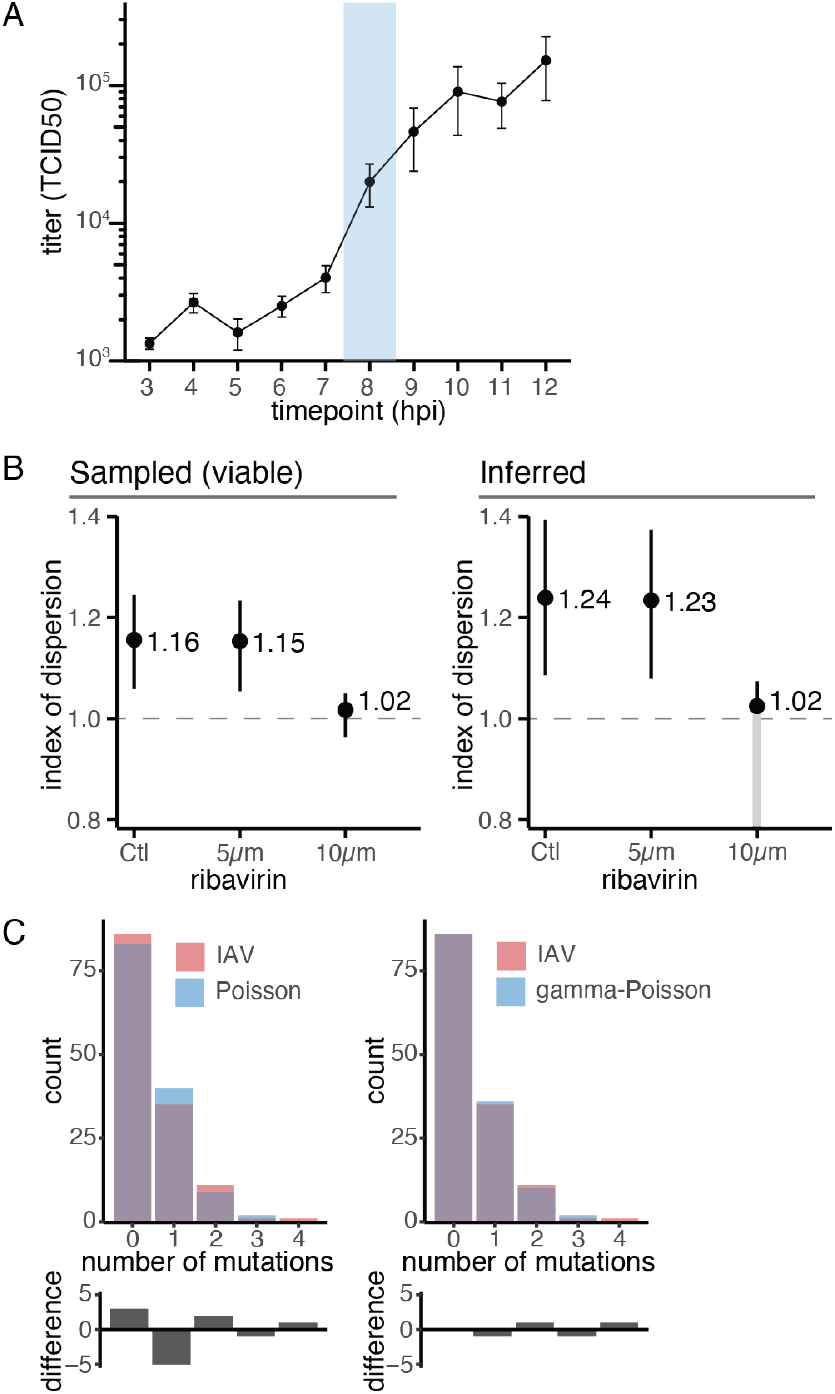
Mutation accumulation for a single round of replication of the H3N2 virus A/Netherlands/499/2017. A) Growth curve of the H3N2 virus A/Netherlands/499/2017 in MDCK-SIAT1 cells (MOI = 0.1). Blue shaded region indicates the timepoint selected for collection of viral supernatants in mutation accumulation experiments. Data represent 3 biological replicates per timepoint, shown are mean +/- standard deviation. B) MDCK-SIAT1 cells were treated with PBS or ribavirin prior to infection (3 hours) and during infection (8 hours, MOI = 0.1). End-point dilutions were deep-sequenced to identify fixed mutations (frequency > 95%). Index of dispersion is the ratio of the variance to the mean of the mutation counts. Left panel: index of dispersion of sampled (viable) genomes, directly measured. Right panel: index of dispersion inferred for all genomes (viable and non-viable), assuming 30% of mutations are lethal (see Methods). Number of sequenced clones per sample after quality control: 134 (Ctl), 96 (5μm), 89 (10μm). Points indicate mean; segments correspond to bootstrap 95% confidence intervals. The lower bound of the 95% confidence interval for the inferred 10μm condition cannot be calculated, as underdispersion is incompatible with a gamma-Poisson distribution (see Methods). C) Comparison of mutation accumulation data (Ctl) to expected counts from a Poisson distribution with the same mean (left), and gamma-Poisson distribution with the same mean and variance (right).

We additionally tested whether treatment with ribavirin, a mutagenic drug, would affect the level of overdispersion. We observed a minimal effect of ribavirin on overdispersion at 5μM drug, and decreased overdispersion at 10μM (Figure 3B, left panel). Given that we tested relatively few clones per treatment, particularly at 10uM ribavirin, it is possible that our observed values are an inaccurate representation of the true level of overdispersion in the viral population. More importantly, our mutation accumulation experiment can only identify mutations in viral clones that are viable, given that genomes with more mutations are more likely to be non-viable, and may under-count the true mutation counts and distribution. However, using Bayes theorem (see Methods), we can infer the mean and variance of the entire viral population, given that ∼30% of random mutations in H3N2 are lethal (20). We found that the inferred level of overdispersion for the whole IAV population is higher than for the viable sample (Figure 3B, right panel). Using the same formula, we calculate the unconditional probability of a progeny virus being viable to be 0.809 for our control sample. Finally, our empirical counts more closely match expected counts from a gamma-Poisson distribution than a Poisson (Figure 3C). In all, given that 1) viral mutation rates evolve, 2) viral mutation rates can be high enough to generate structural and biochemical variability in the polymerase in a single generation, and 3) our empirical evidence of overdispersion in IAV populations, we suggest that viral mutations are more accurately modeled using the gamma-Poisson distribution than the Poisson distribution.

### Implications for lethal mutagenesis of viruses

Lethal mutagenesis is a strategy to achieve extinction of viral populations through artificial elevation of mutation rates (typically via mutagenic drug treatment). As most mutations are deleterious (20–24), elevated mutation rates decrease the number of successful offspring per parent, thus leading to extinction. Under an assumption of Poisson-distributed mutations, the threshold at which extinction occurs (the *extinction threshold*) is a function of the deleterious mutation rate and fecundity--the number of viable offspring per individual (4) (Equation 1 and Figure 4A, black line). The right-hand side of Equation 1 corresponds to the average number of surviving offspring per individual; if this number is greater than one the population survives, if it is less than one the population ultimately goes extinct. The value e^-*U*^ is simply the proportion of offspring with no new mutations (i.e., the null class, according to the Poisson distribution). In this basic model, all mutations are assumed to be deleterious.

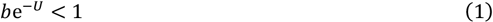

where *b* is fecundity and *U* is the deleterious mutation rate

**Figure 4.**
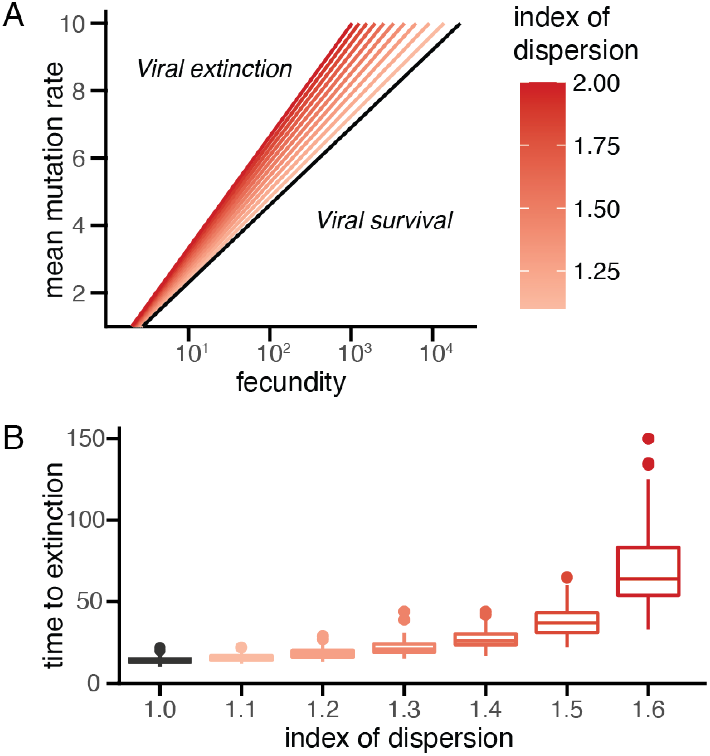
Overdispersion increases the lethal mutagenesis extinction threshold. A) Diagram of the extinction threshold for RNA viruses with diverse mutation rates. Viruses with mutation rate and fecundity that fall below the threshold will survive, while viruses above the threshold will go extinct. The position of the threshold depends upon the index of dispersion, with overdispersion leading to higher extinction thresholds. Mutation rate is the mutations per replicated genome; fecundity is the number of offspring per individual. Extinction thresholds were calculated according to equations 1 and 2, with the Poisson threshold in black, and gamma-Poisson thresholds in red. B) Time to extinction (generations) and index of dispersion for 100 simulated viral populations with fecundity = 10, fitness cost per mutation = 0.4, mean mutation rate = 3, and starting population size = 100. Simulations were performed as described in (23). All simulated populations eventually go extinct according to extinction thresholds given by Equations 1 and 2.

However, we demonstrated above that the assumption of Poisson-distributed mutations may not be valid for viral populations. We can instead derive the extinction threshold based on the null class in a gamma-Poisson model, in which the *mean* mutation rate, the index of dispersion, and the fecundity are parameters (Equation 2, see Methods).

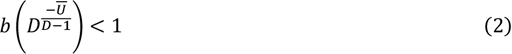

where *D* is the index of dispersion

Using this equation, it becomes clear that overdispersion increases the extinction threshold for a given mean mutation rate and fecundity (Figure 4A, red lines). This observation has profound implications for lethal mutagenesis as a drug strategy; if the index of dispersion cannot be accurately measured for viruses targeted by mutagenic drug treatment, there is substantial risk of underestimating the mean mutation rate required for viral extinction.

Furthermore, the number of generations required for viral extinction to occur in a simulated viral population increases with the index of dispersion (Figure 4B). This behavior is explained by increasing proximity of the modeled viral population to the extinction threshold as overdispersion increases. Viral populations closer to the threshold will have more viable offspring than populations farther from the threshold, despite still reproducing below the rate necessary for replacement (see Equation 2). Therefore, it will take longer for populations to go extinct when they are closer to the extinction threshold. In other words, even if the extinction threshold is successfully crossed through drug treatment, underestimating how far the population is away from the threshold by not considering mutation rate variability could mean that the time to extinction is too long to be clinically relevant.

Inherent to the success of lethal mutagenesis is the ability to accurately measure the parameters that determine the extinction threshold. In our refined model of variable mutation rates, these parameters are mean mutation rate, fecundity (number of viable offspring per individual), and the index of dispersion. While the challenges and methods used to measure the first two parameters have been extensively discussed (28), our innovation of modeling mutation counts with the gamma-Poisson distribution alters how mutation rates can be calculated from experimental data. Specifically, the fluctuation test employed by Luria and Delbruck explicitly assumes a Poisson distribution for calculating the rate of mutation per genome per replication from the number of non-revertant bacterial cultures (1) (Equation 3).

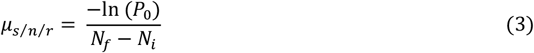

Where *μ*_*s/n/r*_ is mutations per strand per nucleotide per replication, *P*_0_ is proportion of null cultures, and *N*_*f*_ and *N*_*i*_ are the final and initial population sizes, respectively

Since fluctuation tests have since been adapted to measure IAV mutation rates (11, 29), we sought to examine how incorporating the gamma-Poisson distribution would alter existing mutation rate estimates from these assays (Equation 4, see Methods).

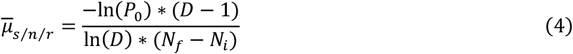

Using our own fluctuation test data from a another H3N2 IAV (A/Hong Kong/4801/2014), we re-calculated the mutation rate using the inferred index of dispersion for our control mutation accumulation sample (11) (Figure 3A). We found that the refined mutation rate estimates are >10% higher than previously calculated (Table 1).

**Table 1.** Mutation rates for the H3N2 strain A/Hong Kong/4801/2014

| **Table 1.** |  |  |
| --- | --- | --- |
| **Mutation class** | **Mutation rate**<br>**(Poisson)** | **Mutation rate**<br>**(gamma-Poisson)** |
| A -> C | $3.38 * 10^{-5}$ | $3.77 * 10^{-5}$ |
| A -> G | $3.02 * 10^{-4}$ | $3.37 * 10^{-4}$ |
| A -> U | $1.31 * 10^{-5}$ | $1.47 * 10^{-5}$ |
| C -> A | $1.73 * 10^{-5}$ | $1.93 * 10^{-5}$ |
| C -> G | $9.74 * 10^{-6}$ | $1.09 * 10^{-5}$ |
| C -> U | $4.59 * 10^{-5}$ | $5.12 * 10^{-5}$ |
| G -> A | $7.22 * 10^{-5}$ | $8.05 * 10^{-5}$ |
| G -> C | $2.77 * 10^{-5}$ | $3.09 * 10^{-5}$ |
| G -> U | $6.01 * 10^{-5}$ | $6.71 * 10^{-5}$ |
| U -> A | $4.54 * 10^{-6}$ | $5.07 * 10^{-6}$ |
| U -> C | $3.10 * 10^{-4}$ | $3.48 * 10^{-4}$ |
| U -> G | $3.65 * 10^{-5}$ | $4.07 * 10^{-5}$ |
| **Overall** | **$2.51 * 10^{-4}$** | **$2.81 * 10^{-4}$** |

This finding, combined with the elevation of extinction thresholds due to overdispersion, further highlights the impact that modeling mutations with the gamma-Poisson distribution has on predictions about the success of lethal mutagenesis as an antiviral drug strategy.

## Discussion

We have presented empirical evidence that mutation rates in IAV populations are variable, and thus the number of mutations per offspring will follow a distribution that accounts for rate variability, such as the gamma-Poisson. It is likely that mutation rate variability is common among other RNA viruses, given that most have similarly high mutation rates. Furthermore, mutation rate variability (and thus gamma-Poisson distributed mutation) was also recently observed in a yeast model of cancer (18), and may be a hallmark of rapidly evolving populations in general. Our finding of viral mutation rate variability has direct implications for lethal mutagenesis, an important antiviral drug strategy. Specifically, addressing mutation rate variability in models of lethal mutagenesis significantly increases the extinction threshold required for treatment success. The consequences of underestimating the extinction threshold when using mutagenic drugs can be significant, as this leads to an increase in mutations without causing extinction of the viral population. Recent studies of SARS-CoV-2 viruses from patients treated with the mutagenic drug molnupiravir have identified increased mutations that gave rise to persistent mutagenic signatures among downstream cases (9). This observation can be explained by failure of the molnupiravir treatment to push SARS-CoV-2 populations beyond the true extinction threshold.

Accurate estimation of the parameters that determine the extinction threshold is critical to the success of lethal mutagenesis as an antiviral drug strategy. Not only must mutation rate and viral burst size be measured, as previously understood (28), but, given our new model of mutation rate variability, the index of dispersion must also be determined. In this work we demonstrate that mutation accumulation experiments can be used to empirically estimate this parameter, if enough clones are characterized. While performing infections in a controlled cell culture environment allowed us to focus on the genetic contribution to mutation rate variability, we could not completely control for environmental and stochastic causes. However, our conclusions – that viral mutation rates are more accurately modeled with the gamma-Poisson distribution and that overdispersion increases the extinction threshold – hold true regardless of the source of variability.

We also found that accounting for overdispersion increases the IAV mutation rate as estimated in Luria-Delbrück fluctuation tests. It is less clear how the index of dispersion may change as mutation rates are elevated through mutagenic drug treatment. While our current data examining the effect of ribavirin treatment on the index of dispersion is inconclusive due to low sample sizes, it is crucial for future experiments to determine if there is a relationship between mutation rate and index of dispersion. Previous theoretical work has shown that the mean number of mutations increases with mutation rate variance among loci with sufficient positive epistasis(25). According to our lethal mutagenesis model, a positive relationship between mutation rate and variance would result in the extinction threshold shifting ever higher upon treatment with mutagenic drugs, further increasing the risk that the threshold is not reached.

While we have used our improved understanding of viral mutation rates to refine the most basic model of lethal mutagenesis, we have not explored how more complex models would change with a gamma-Poisson model of mutation accumulation. Specifically, the basic model used in this work assumes that all mutations are equally deleterious and does not consider heritability of mutation rates. Previous theoretical work has shown that assuming perfect heritability would increase the number of mutations per genome and thus decrease population fitness, but selection will ultimately limit heritability (25). As most mutations in the IAV polymerase are deleterious, the effect of heritability on the extinction threshold can be assumed to be minimal (20, 26, 27). It will be important for future work to consider the effect of mutation rate variability on extinction thresholds in the context of heritability and more complex fitness landscapes. Each increase in model complexity can yield losses in interpretability, yet a full picture of the forces driving viral extinction during lethal mutagenesis is critical to ensure positive public health outcomes from mutagenic drug treatment.

## Materials and Methods

### Availability of data and computer code

Illumina sequencing data are available in the Sequencing Read Archive (SRA) at accession PRJNA1259473. Source code used to analyze the data, perform all modeling and statistical analysis, and generate figures is available at https://github.com/lauringlab/MutationRateVariability.

### Viruses, plasmids, and cells

MDCK-SIAT1 cells and HEK-293T cells were maintained in Dulbecco’s modified Eagle medium (Gibco 11965) supplemented with 10% FBS (Gibco 10437), 25mM HEPES (Gibco 15630), and 0.1875% BSA fraction V (Gibco 15260). Plasmids encoding each 8 segments of the H3N2 strain A/Netherlands/499/2017 were provided by Dr. Ron A. Fouchier (Erasmus University Medical Center, Rotterdam, Netherlands). The virus was rescued in 6-well plates by transfection of a co-culture of 293T (3 x 10^5^ cells/well) and MDCK-SIAT1 (1.5 x 10^5^ cells/well) using 1μg of each plasmid and 2μL of TransIT-LT1 (Mirus 2300) per μg of DNA in Opti-MEM (Gibco 31985) (Hoffmann 2000). After 24 hours, the media were changed to viral infection media (Dulbecco’s modified Eagle medium [Gibco 11965] supplemented with 25mM HEPES [Gibco 15630], 0.1875% BSA fraction V [Gibco 15260], and 2ug/mL TPCK-trypsin). Viral supernatants were harvested 72 hours after transfection with 0.5% glycerol and clarified by centrifugation prior to snap-freezing. Transfection titers were determined by tissue culture infective dose (TCID50) assay. Passage 1 stocks were made by infecting T25 flasks containing 8 x 10^5^ MDCK-SIAT1 cells at MOI = 0.01 in 1mL of viral infection media. Infections were rocked every 15 minutes for 1 hour, then the media were aspirated and replaced with 3mL fresh viral infection media. Infection supernatants were collected after 48 hours and titered by TCID50.

### Growth curve

24 hours prior to the experiment, 6.5 x 10^4^ MDCK-SIAT1 cells/well were plated in 24-well plates. On the day of the experiment, cells were infected with passage 1 stocks of A/Netherlands/499/2017 at MOI = 0.1 in 120uL viral infection media for 1 hour, rocking every 15 minutes. Media were then replaced with 500uL fresh viral infection media. Viral supernatants were then collected each hour from 3 to 12 hours post infection and then titered by TCID50 in duplicate.

### Mutation accumulation

Twenty-four hours prior to the experiment, MDCK-SIAT1 cells were plated in 6-well plates at a density of 2.5 x 10^5^ cells/well. Three hours prior to infection, cells were treated with 0um, 5uM, or 10uM ribavirin (Sigma-Aldritch R9644). After 3 hours, cells were infected with passage 1 stocks of A/Netherlands/499/2017 at MOI = 0.1 in 500uL viral infection media with 0uM, 5uM, or 10uM ribavirin for 1 hour, rocking every 15 minutes. Media were then replaced with 2mL fresh viral media with 0uM, 5uM, or 10uM ribavirin. Viral supernatants were collected 8 hours after infection and titered by TCID50. These titered supernatants were diluted to generate a ∼20% chance of containing a single infectious unit per 100uL of viral infection media (∼4 TCID50/mL) and 100uL of these dilutions were then plated with 6000 MDCK-SIAT1 cells per well in 96-well plates and incubated until complete cytopathic effect (CPE), ∼5 days. Supernatants from CPE+ wells were then collected, and RNA was harvested for Illumina sequencing.

### Illumina library preparation, sequencing, and data analysis

Nucleic acid was extracted from clarified viral supernatants using the MagMAX Viral/Pathogen II Nucleic Acid Isolation Kit on a KingFisher instrument. The 8 genomic segments of IAV were amplified in a one-step multiplex RT-PCR (30). Amplified cDNA were prepared for sequencing using the Illumina DNA Prep M Tagmentation Kit and sequenced on a NextSeq instrument with 2x150 PE reads. Sequence reads were trimmed using cutadapt (Version 1.18) (31) and aligned to the A/Netherlands/499/2017 genome sequence using BWA (version 0.7.17) (32). Duplicate reads were removed using picard (version 2.8.1) (33) and single-nucleotide variants (SNV) were called using iVar (34); prior to variant calling, reads were filtered to have a mapping quality ≥20, a phred score ≥30. Identified SNV were then filtered to require a per-site sequencing depth of ≥100. To account for mutations in the original viral stock, we searched for any mutations reaching 100% frequency in a majority of samples. Only 1 mutation matched these criteria (PB2 A1700G) and was masked from downstream analysis. Next, we identified samples in which multiple viral clones may have initiated an infection in that well by filtering for samples with mutation frequencies between 15% and 95%. Our rationale is that if an infection was truly seeded with a single clone, then mutations should either be close to 100% (due to mutations in the infecting clone) or very low frequency (due to spontaneous mutation as the infection proceeded). After this filtering step, we were left with 134 (Control), 96 (5uM ribavirin), and 89 (10uM ribavirin) analyzed clones. Fixed mutations (>95% frequency) from each clone were then counted.

### Inference of mean and variance of mutation counts in viable and non-viable viruses

The mean and variance of mutation counts in viable and non-viable viruses was inferred using Bayes theorem. Briefly, our data reflect the distribution of mutation counts conditional upon viral viability, *P*(*X* = *m* | *viable*). Therefore, the equation for the unconditional distribution of mutation counts is:

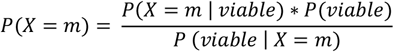

The probability that a virus is viable given the number of mutations is a function of the probability that a random mutation is lethal, *P*_*L*_:

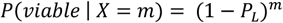

And as *P*(*viable*) does not depend upon the value of *X*, and *P*(*X* = *m*) must sum to 1, we can calculate the unconditional distribution of mutation counts using:

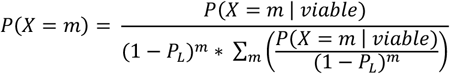

Finally, the value of *P*_*L*_ has been previously estimated to be ∼30% (20).

### Derivation of gamma-Poisson extinction threshold and Luria-Delbrück mutation rate

The lethal mutagenesis extinction threshold can be derived starting with the gamma-Poisson distribution using parameters for mean deleterious mutation rate (*Ū*) and index of dispersion (*D*):

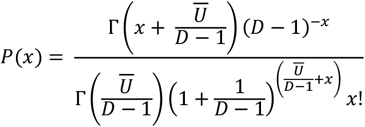

Then, the equilibrium mean fitness is simply the density at *x* = 0:

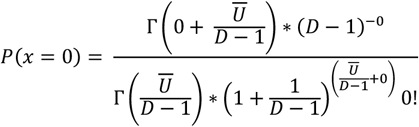

Which simplifies to:

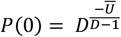

The extinction threshold is then:

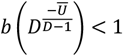

The mutation rate from a Luria-Delbrück fluctuation test can be calculated by first solving the gamma-Poisson density function at *x* = 0 for the mean mutation rate, *Ū*:

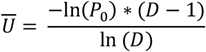

And then normalizing by population growth:

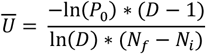

## Acknowledgments

We thank Will Fitzsimmons and Chris Blair for their assistance in Illumina library preparation, and Doctor Emily Bendall for her help with sequencing data analysis. We also thank Doctors James J Bull, Luis Zaman, and Jonathan Terhorst for their helpful discussions regarding lethal mutagenesis theory and evolutionary simulation. This work was supported by National Institute of Allergy and Infectious Diseases R01AI170520 and a Burroughs Wellcome Fund Investigator in the Pathogenesis of Infectious Diseases Award (both to Adam S Lauring).

